# Chronic cortisol exposure in early development leads to neuroendocrine dysregulation in adulthood

**DOI:** 10.1101/856633

**Authors:** Ellen I. Hartig, Shusen Zhu, Benjamin L. King, James A. Coffman

## Abstract

**Objective:** Chronic early life stress can affect development of the neuroendocrine stress system, leading to its persistent dysregulation and consequently increased disease risk in adulthood. One contributing factor is thought to be epigenetic programming in response to chronic cortisol exposure during early development. We have previously shown that zebrafish embryos treated chronically with cortisol develop into adults with constitutively elevated whole body cortisol and aberrant immune gene expression. The objective of the experiments reported here was to further characterize the phenotype of those adults.

**Results:** We find that adult zebrafish derived from cortisol-treated embryos have aberrant cortisol levels, tissue distribution, and dynamics, which correlate with differential activity of key glucocorticoid-responsive regulatory genes *klf9* and *fkbp5* in blood and brain.

## Introduction

Epidemiological studies have shown that early life stress (ELS) can increase later life risk of developing a variety of physical and mental health problems, many of which are linked to chronic inflammation [1–3]. This long-term ‘developmental programming’ has been hypothesized to in part be an epigenetic effect of chronic exposure to elevated cortisol levels [4, 5]. Consistent with this, chronic glucocorticoid (GC) exposure, such as occurs with chronic stress, genetic diseases such as Cushing’s syndrome, or with extended GC therapy, promotes development of metabolic and inflammatory disease [6]. It is thought that developmental programming in response to ELS can be adaptive to the extent that it functions in an adaptive and anticipatory fashion to tune the responsiveness of the stress system to the stressfulness of the environment [1, 7]. However, stress-induced programming can be maladaptive when there is a mismatch between the adult environment and that which was encountered during earlier development, as the programming may result in levels of GC reactivity and/or resistance inappropriate to the level of stress actually encountered in adulthood [1]. Moreover, the cost of such programming can be an increased allostatic load that promotes unhealthy aging [8].

Zebrafish are emerging as an excellent model system for experimental studies of stress-induced developmental programming [9–13]. We recently showed that zebrafish embryos treated chronically with 1 µM cortisol develop into adults that maintain elevated whole body cortisol levels and aberrantly express pro-inflammatory genes, with higher basal expression levels in peripheral tissues but a blunted response to tailfin injury or lipopolysaccharide injection [13], indicating that chronic GC exposure during early development has long-term effects on the neuroendocrine stress axis and GC-regulated gene expression. Those results are extended by the experiments reported here.

## Main Text

### Results

#### Adults derived from cortisol-treated embryos display aberrant cortisol tissue distribution and dynamics

Zebrafish embryos were treated chronically with cortisol or vehicle for the first five days of development as described previously [13], then raised to adulthood (5+ months), at which time cortisol levels in different tissues were measured in both fed and fasted fish (Fig. 1A, B). This analysis showed that the higher whole-body cortisol levels that we previously observed in the treated fish (which had been fed) are largely due to higher levels in kidney (which when dissected out likely contains the cortisol-producing interrenal gland) and blood, whereas levels in brain were actually somewhat lower in the treated group (Fig. 1B). However, in fish that had been fasted for 24 hours, cortisol levels were lower in blood of the cortisol-treated group compared to controls, but higher in the brain, as well as skin and gut (Fig. 1B). Plotting of the ratios of cortisol levels in fed and fasted fish reveals that while 24 hours of fasting did not affect brain cortisol levels in control fish, it caused brain cortisol levels to quadruple in the treated fish (Fig. 1B, right panel). Compared to their control siblings, fasted adults derived from cortisol-treated embryos consistently displayed lower blood cortisol with a compressed dynamic range (Fig. 1B-D), possibly owing to the concomitant spike in brain cortisol, which would be expected to downregulate the hypothalamus-pituitary-adrenal/interrenal axis. Consistent with this, in fasted fish derived from cortisol-treated embryos, blood cortisol levels were similar to those of their untreated siblings exposed to Dexamethasone (Dex) for 8 hours, and were not further reduced by the Dex treatment (Fig. 1D). Moreover, brain expression of the ACTH-encoding gene *pomca*, a target of glucocorticoid receptor (GR)-mediated repression, is significantly lower in fasted fish derived from cortisol-treated embryos, and is less affected by Dex (Fig. 1E). These data show that on the whole cortisol is both chronically elevated and aberrantly regulated in adults derived from cortisol-treated embryos, with a compressed dynamic range in blood that correlates with an expanded dynamic range in brain and peripheral tissues compared to controls.

**Figure 1.**
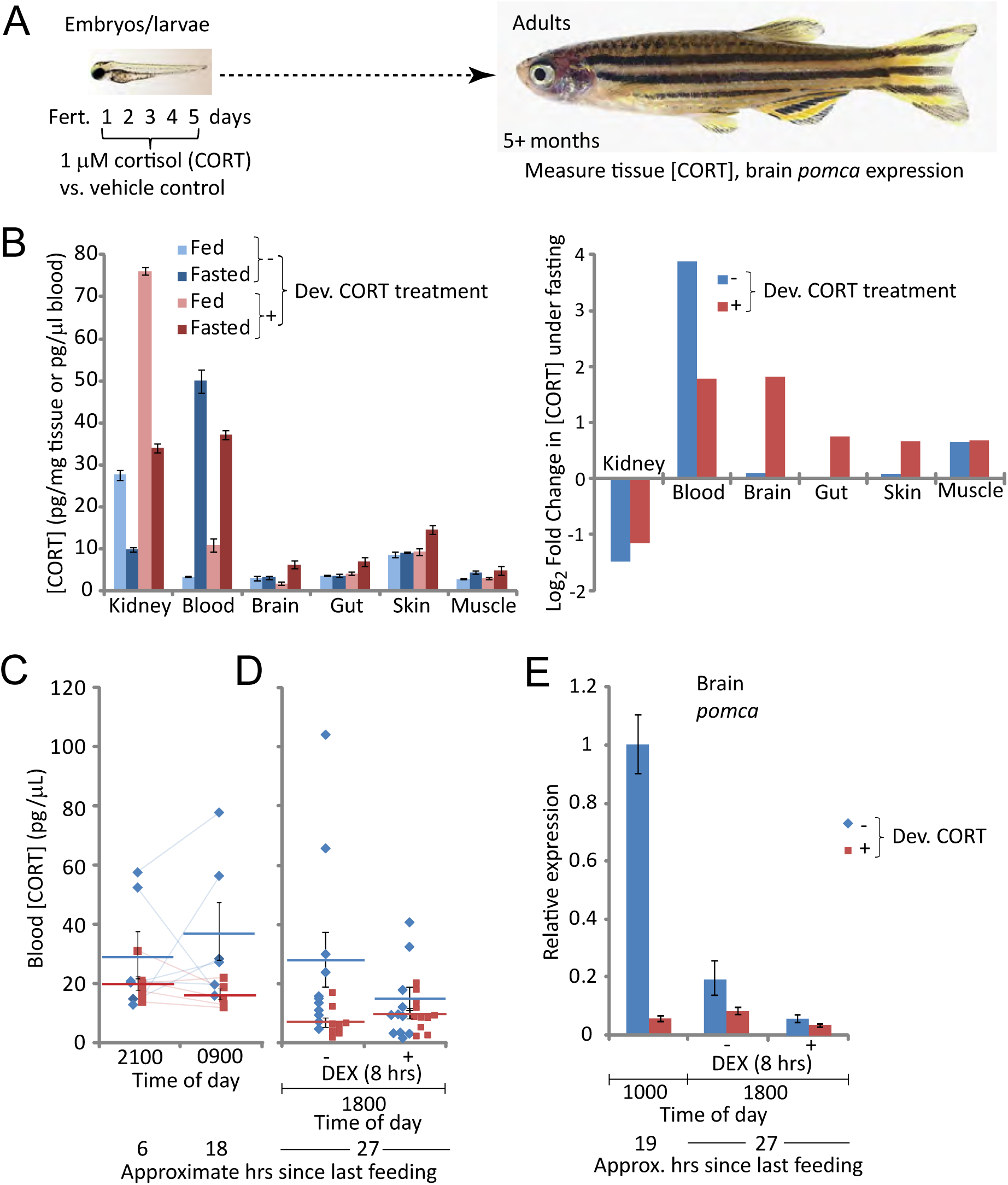
Embryos treated chronically with cortisol develop into adults with aberrant cortisol levels, tissue distribution, and dynamics. (A) Schematic of experimental design; (B) Cortisol levels in different tissues of fed and fasted (24 hours) fish (left panel) and the inferred change in those levels induced by fasting (right panel). Each measurement was taken from pooled tissues of 6 fish, with equal numbers of males and females. Error bars are the standard deviations of technical replicates. (C) Cortisol levels in blood draws from single fish at night and the following morning, with the amount of time since the last feeding indicated. Thin lines between data points indicate repeat measurements from the same fish. Averages +/− standard errors of the mean (SEM) are also shown. (D) Fasting cortisol levels in blood of individual fish (data points) following 8 hours exposure to 1 µM dexamethasone (DEX) or vehicle. The averages +/− SEM for each group are also indicated. (E) Relative expression of *pomca* in brain tissue in the late morning, as well as later the same day following 8 hours exposure to DEX or vehicle. The bars indicate averages +/− the standard deviation of 3 qPCR readings (technical replicates). For each measurement brain tissue was pooled from 5-6 fish, mixed males and females in equivalent proportions within each comparison group (controls 1 male and 4-5 females; cortisol-treated 2 males and 3 females).

#### Adults derived from cortisol-treated embryos differentially express the glucocorticoid-responsive regulatory genes klf9 and fkbp5 in blood and brain

To identify genes persistently affected in adults derived from cortisol-treated embryos at the level of chromatin we performed ATAC-seq [14, 15] on blood cells isolated from three independent replicates comparing ~1-year old adults derived from cortisol-treated embryos and their vehicle-treated (control) siblings. Each ATAC-seq peak was scored according to its size in cortisol-treated fish relative to controls, with higher scores indicating proportionally more sequence reads and hence chromatin with a more open configuration in the cortisol treated group. Of ~20,000 scored peaks, that with the 3^rd^ highest score encompassed the promoter of *klf9* (Fig. 2A; Table 1), a known GR target gene that functions as a feedforward regulator of GR signaling [16–20]. Several additional genes with high peak scores are also known GR targets in mammals, including *fkbp5* (Fig. 2A; Table 1) a clinically important feedback regulator of the GR [17, 21–24]. Three of the genes in the top 35 (*chac1*, *klf9*, and *fkbp5*) were found in our previous study to be highly upregulated in larvae chronically exposed to cortisol (Table 1). Interestingly, in a HOMER motif enrichment analysis [25] of the 251 peaks with scores ±100, four of the top ten scoring motifs were consensus binding sequences for krüppel-like factors, including Klf9 (Supp. Fig. S1).

**Figure 2.**
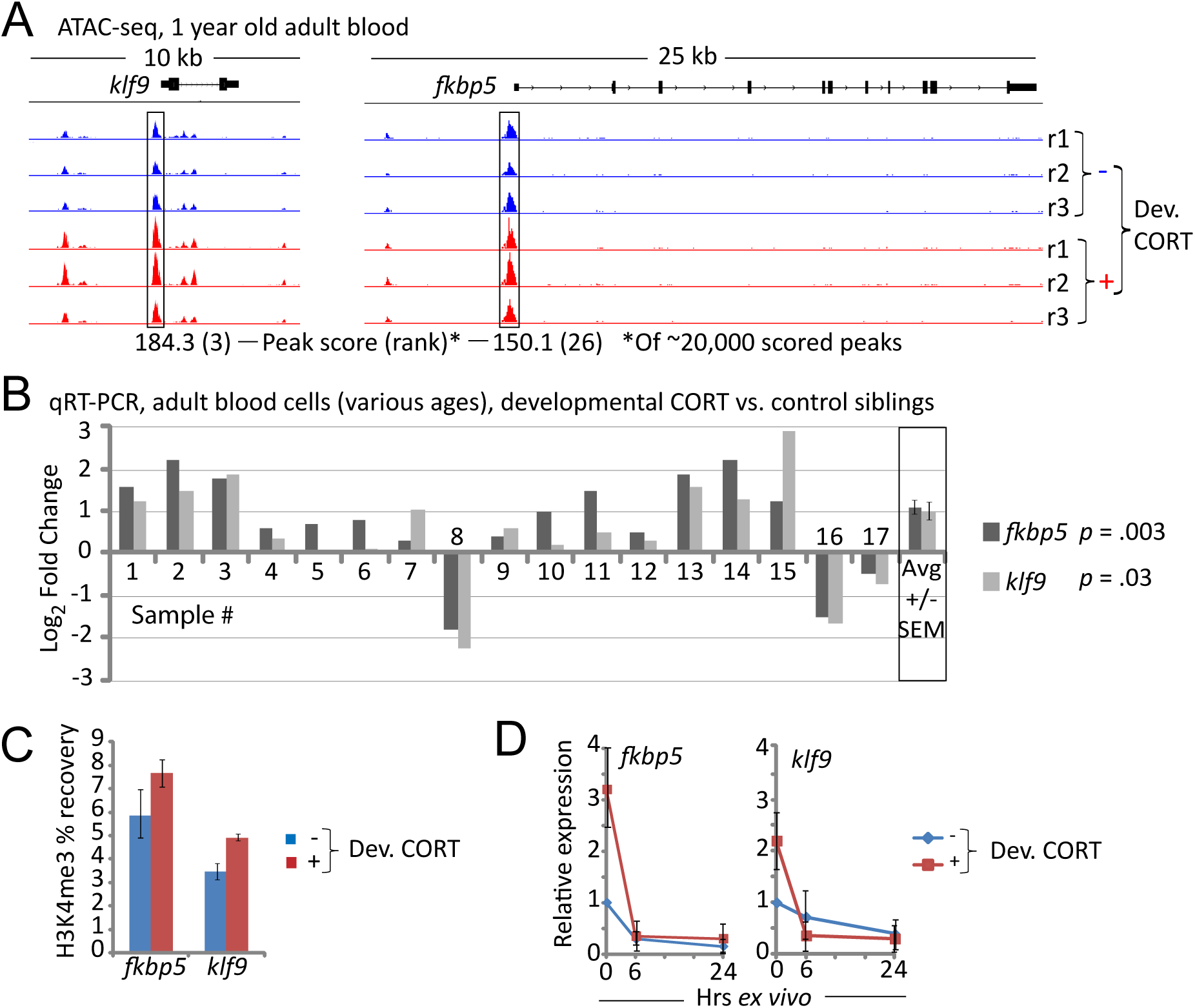
The regulatory genes *klf9* and *fkbp5* have higher on average activity in blood cells of adults derived from cortisol-treated embryos. (A) ATAC-seq peaks associated with *klf9* and *fkbp5*, from three biological replicates of each treatment (r1-r3). For each replicate sample blood was pooled from 6 individuals mixed sex with equal representation of males and females. (B) Relative expression of *klf9* and *fkbp5* in 17 samples of adults from different experimental cohorts of cortisol-treated embryos compared to their control siblings. The averages +/− SEM are also shown. For each experimental sample blood was pooled from 6 individuals mixed sex with equal representation of males and females. (C) ChIP-qPCR of H3K4me3 levels in the promoter regions of *klf9* and *fkbp5* from a single sample of blood pooled from 6 individuals (mixed sex, equal representation). (D) Relative expression of *klf9* and *fkbp5* in blood cells immediately after being drawn, and then after 6 or 24 hours of *ex vivo* culture in the absence of cortisol. The plots represent the grand means +/− the SEM of three biological replicates, each done on blood samples pooled from 6 individuals of mixed sex.

**Table 1:**
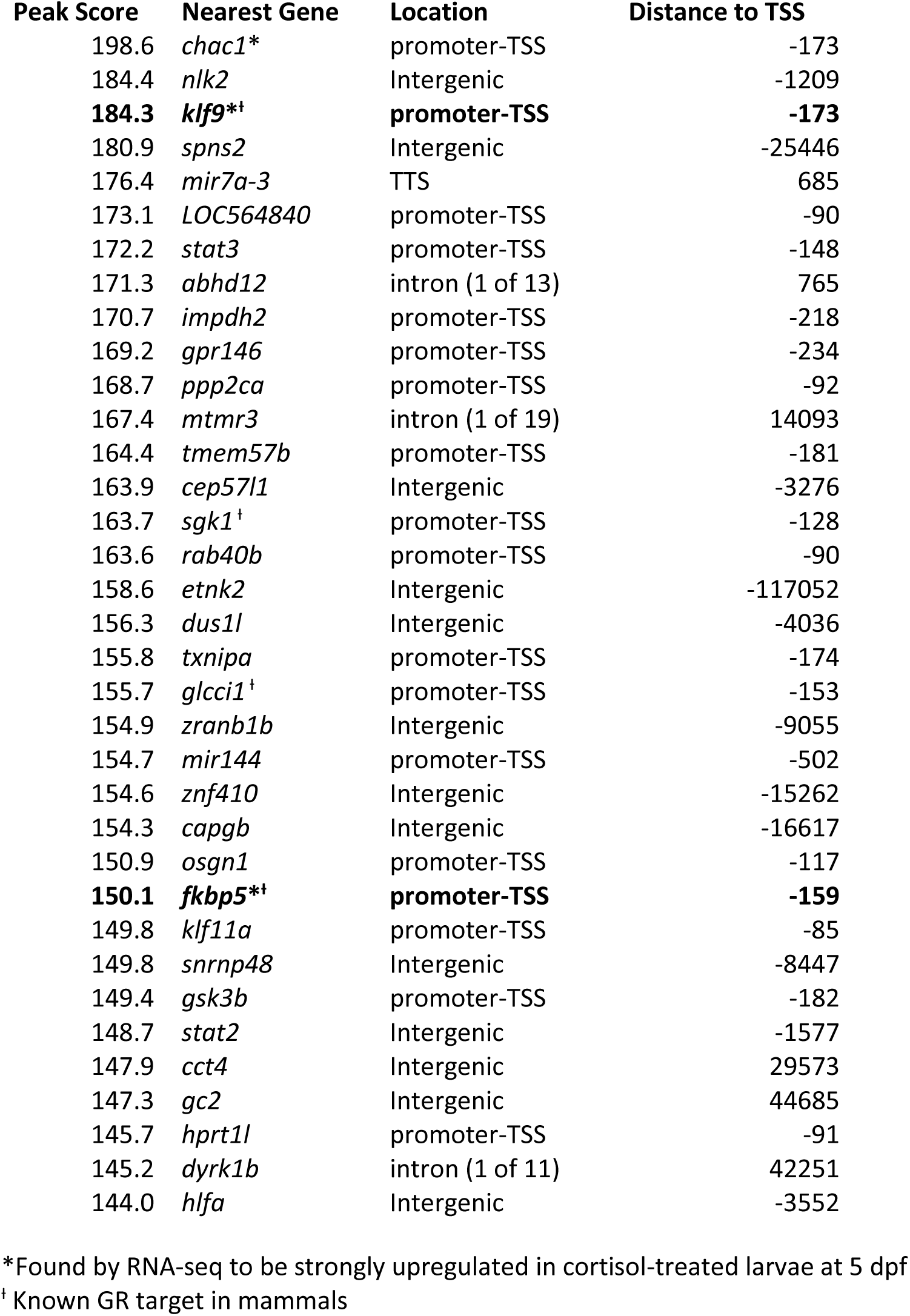
Top 35 ATAC-seq peaks in blood cells of 1-year old fish derived from cortisol-treated embryos.

The ATAC-seq results suggest that in adults derived from cortisol-treated embryos the persistence of chronically elevated cortisol levels upregulates the activities of both *klf9* and *fkbp5*. To further test this, quantitative reverse transcription and polymerase chain reaction (qRT-PCR) of both genes was carried out on blood cells of adults derived from cortisol-treated embryos and their vehicle-treated siblings. In 14 out of 17 independent blood cell samples, *klf9* and/or *fkbp5* expression was found to be elevated in adults derived from cortisol-treated embryos compared to their control siblings, both genes being overexpressed ~2-fold on average for all samples (Fig. 2B). The fact that *fkbp5* and *klf9* expression was elevated in most, but not all the blood samples is consistent with the observation that blood cortisol in the treated group had a compressed dynamic range compared to controls (Fig. 1B, right panel). Chromatin immunoprecipitation detected higher levels of H3K4 trimethylation in the promoter regions of both genes in the treated fish (Fig. 2C), suggesting that the elevated expression observed in blood cells of those fish is transcriptional. This is likely a direct effect of elevated blood cortisol levels, as, when blood cells were cultured *ex vivo* the expression of both genes decreased over time and converged (Fig. 2D).

In an initial experiment that surveyed the expression of 95 candidate genes in adult brains, *fkpb5* and *klf9* stood out as the two that were most highly overexpressed in the cortisol-treated group compared to their control siblings (Supp. Fig. S2A). However, this result was not reproduced in subsequent measurements from additional experimental cohorts (Supp. Fig. S2B-F), suggesting that the fish used for the initial survey may have recently experienced a stressor that produced a spike in brain cortisol levels of the treated group (e.g., as observed with fasting; Fig. 1B). In any case, one strikingly consistent finding from the gene expression measurements made from both blood cells (Fig. 2B) and brain (Supp. Fig. S2) was that expression of *fkbp5* and *klf9* was highly correlated: when one of the genes was overexpressed in the treated group, so was the other, and *vice versa*. Finally, although most of the gene expression measurements were made from pools of mixed sexes containing equal numbers of matched-sized males and females, in an experiment where the sexes were segregated we found that for both genes brain expression levels were higher in males, and that the differential expression in treated and control groups was greater in males than in females (Supp. Fig. S2F), indicating that activities of both genes are also regulated by sex.

### Conclusions

These results extend our earlier findings showing that in zebrafish, chronic glucocorticoid exposure during early development leads to persistent dysregulation of the neuroendocrine stress system and associated gene expression in adulthood. The data indicate that the neuroendocrine dysregulation occurs a multiple levels, including cortisol production (higher in treated fish), delivery of cortisol from circulation to tissues (lower blood retention in the treated fish after fasting, with concomitantly higher levels in brain and peripheral tissues), and activity of key genes that regulate GC signaling dynamics in receiving cells (*klf9* and *fkbp5*). Since similar phenotypes are associated with ELS and chronic glucocorticoid exposure in humans, our results underscore the relevance of zebrafish as a model organism for investigating the molecular-genetic and cellular mechanisms underlying this important public health issue.

### Methods

#### Zebrafish procedures

Wild-type zebrafish of the AB strain maintained in the MDI Biological Laboratory zebrafish facility as previously described [13] were used for all of the reported experiments. Cortisol treatment of larvae was carried out as previously described [13]. For the dexamethasone treatments adult zebrafish were removed from the re-circulating system and transferred into 1 L holding tanks containing either 1 µM dexamethasone or 2.5 ppm vehicle (DMSO) in system water. For tissue collection fish were euthanized in tricaine as previously described [13].

#### Ex Vivo Blood Cell Culture

Blood was collected by a nonlethal venous puncture under the operculum. 1µL samples of blood from 6 fish of mixed sex per condition were pooled in heparin (5 I.U./mL) in PBS, washed, and plated at a density of 1 million cells/mL in supplemented L-15 media (L-glutamine, 10% FBS, 25 mM HEPES, 2.2 g/L sodium bicarbonate, Primocin). Cells were cultured at 28° C in 5% CO_2_.

#### Cortisol measurements

Cortisol was measured by ELISA as previously described [13], using dissected tissues or blood samples. In some cases blood was collected by nonlethal venous puncture under the operculum, allowing repeated measurements (e.g. as shown in Fig. 1C).

#### ATAC-seq

ATAC-seq libraries from blood cells (~200,000 cells per sample) from three biological replicates of adults derived from cortisol-treated embryos and their control (vehicle-treated) siblings (10, 11, and 12 months old) were constructed using the Nextera DNA Library Prep kit (Illumina Cat #FC-121-1030), following the procedures provided with the kit with minor modifications (detailed protocol available upon request) [14, 15]. Libraries were sequenced at the Jackson Laboratory NextGen Sequencing Facility (Bar Harbor Maine) using an Illumina NextSeq 500 (Illumina, San Diego, CA). Following initial diagnostic sequence analyses using FastQC (v0.11.8; https://www.bioinformatics.babraham.ac.uk/projects/fastqc/), reads were trimmed using Trimmomatic (v0.32) [27] with parameters ‘ILLUMINACLIP:TruSeq3-PE.fa:2:30:10 LEADING:3 TRAILING:3 SLIDINGWINDOW:4:15 MINLEN:36’. The remaining paired reads were truncated to 30bp reads using fastx_trimmer from the FASTX Toolkit (v0.0.13; http://hannonlab.cshl.edu/fastx_toolkit/) with parameters ‘-Q33 -l 30’. Truncated reads were then mapped to the zebrafish GRCz10 genome assembly (RefSeq assembly accession: GCF_000002035.5) using bowtie (v1.0.0) [28] with parameters ‘--chunkmds 520 -y -v 2 --best --strata -m 3 -k 1 -S -p 8’. Read alignments to the mitochondrial genome were filtered out from the SAM files and the resulting SAM files were converted to BAM files using samtools (v0.1.18) [29]. Nuclease accessible peaks were called using the callpeak function in MACS2 (v2.1.1.20160309) [30] with parameters ‘-f BAM -g 146444456 nomodel -nolabbda -keep-dup all -call-summits -q 0.001 –B’ and Ensembl (v85) [31] annotation of GRCz10. Differential binding events were called using the bdgdiff function in MACS2 with parameters ‘C’ [32]. Motifs within peaks were predicted and annotated using HOMER (v4.8; http://homer.ucsd.edu/home) [25].

#### RNA isolation and quantitative reverse transcription –polymerase chain reaction qRT-PCR

Total RNA was isolated from blood cells or dissected brain tissue using Trizol and ethanol precipitation. qRT-PCR was performed and the data quantified by the ddCt method as previously described [13].

#### Chromatin immunoprecipitation (ChIP)

ChIP was performed on formaldehyde-crosslinked chromatin from blood cells, essentially using the procedure described by Lindeman et al. [33] an anti-H3K4me3 antibody (Abcam Cat. No. Ab8580) (detailed ChIP protocol provided on request). Input and immunoprecipitated DNA samples were analyzed by qPCR with Sybr Green Fastmix. The primers were targeted to promoter regions of *klf9* (CACATCGACACCGCCTTCATCTG; ATCTTGAACCTCGCCGCTGATTG) and *fkbp5* (ACACAACGGGTTACGGGTC; AAATTAAGGGCAGGCCTTGGA). A standard curve was made using serial dilutions of control input DNA and regression analysis was used to calculate percent recovery (immunoprecipitated DNA/ input DNA) × 100%.

## Limitations

- The measurements shown in Figure 1 were done only once.
- When interpreting the single time point measurements reported here it is important to consider that cortisol levels are dynamic, with circadian rhythms as well as ultradian fluctuations with an order of magnitude dynamic range.

## List of Abbreviations

ELS: Early life stress
GC: Glucocorticoid
GR: Glucocorticoid receptor
ATAC-seq: Assay for transposase accessible chromatin - sequencing
qRT-PCR: Quantitative reverse transcription and polymerase chain reaction
ELISA: Enzyme linked immunosorbant assay

## Declarations

### Ethics approval and consent to participate

All animal procedures were approved by the Institutional Animal Use and Care Committee of the MDI Biological Laboratory. This research did not involve human subjects.

### Consent for publication

Not applicable.

### Availability of data and materials

The ATAC-seq dataset generated during this study is available in the NCBI Gene Expression Omnibus (GEO) repository, under accession number GSE137987.

### Authors’ contributions

EIH designed the fed versus fasting cortisol experiment, carried out all of the zebrafish manipulations, tissue dissections, blood draws, and cortisol measurements, as well as qRT-PCR and ChIP analyses from blood.

SZ constructed the ATAC-seq libraries, developed the ChIP protocol that was used in the lab and carried out the qRT-PCR measurements from brain tissue.

BLK carried out the ATAC-seq data analyses.

JAC conceived and designed the overall project, drafted the figures, and wrote the manuscript.

All authors read and approved the final manuscript prior to submission.

## Acknowledgements

Not applicable.

---

**Supplemental Figure S1.**
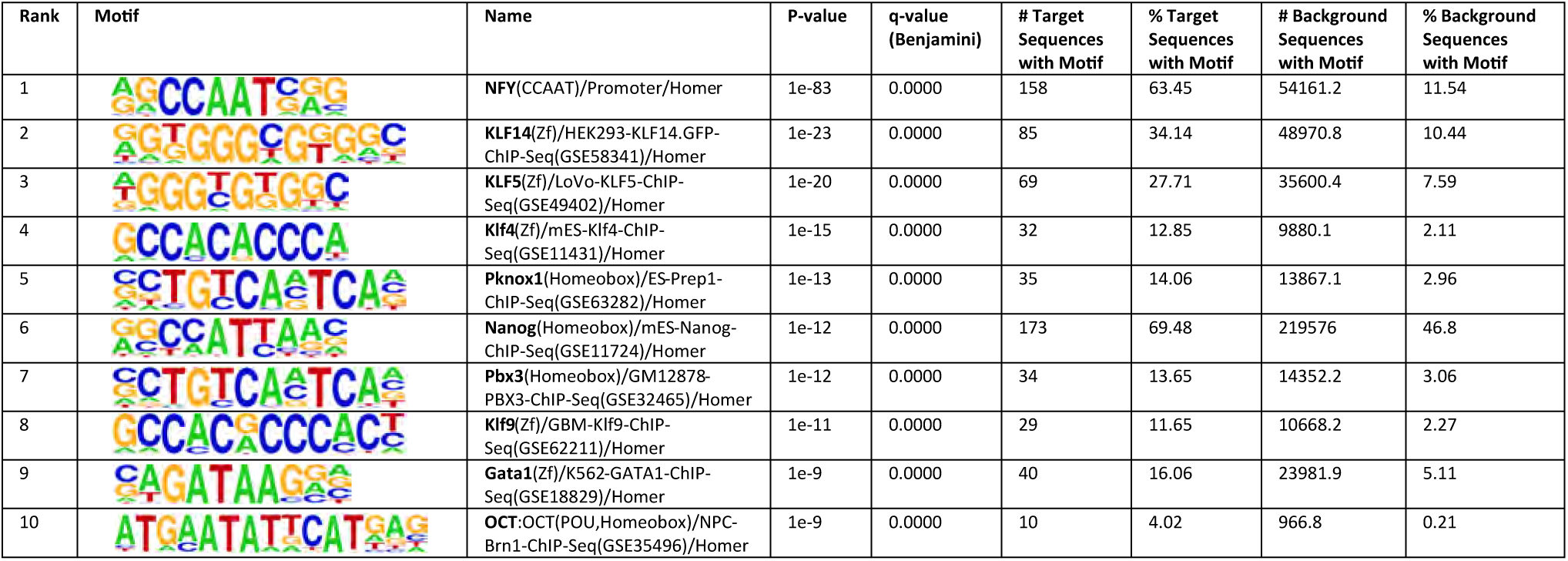
Top ten transcription factor binding motifs identified by HOMER motif enrichment analysis of sequences from the 251 peaks that scored ≥100.

**Supplemental Figure S2.**
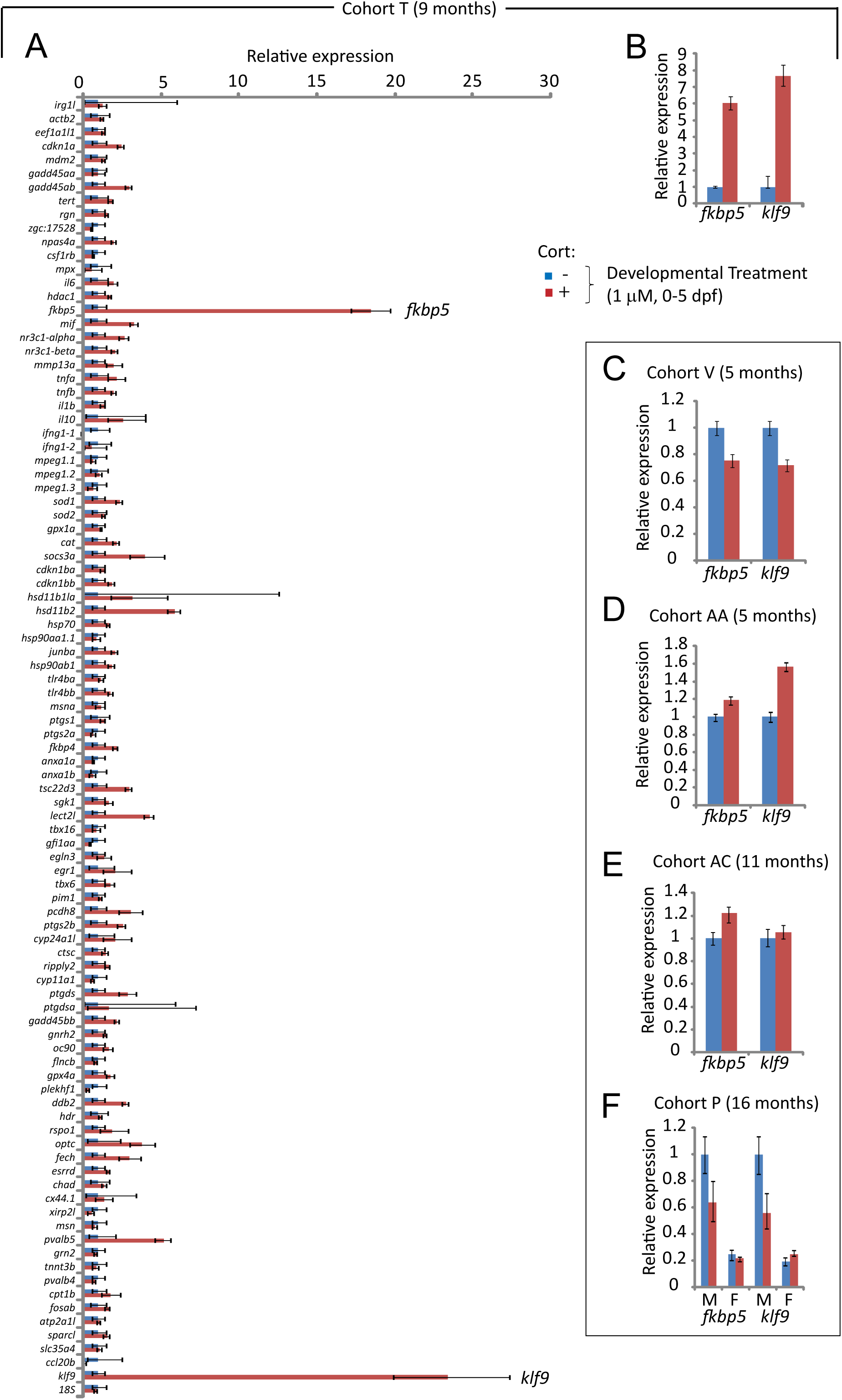
Survey of gene expression by 96-well qPCR array and relative expression levels of *klf9* and *fkbp5* in brain tissue dissected from different experimental cohorts. Each measurement was made from pooled brain RNA of 6 individuals of mixed sex (equal representation), except where indicated (panel F). The qPCR measurement shown in panel (B) is from the same RNA samples that were used to generate the data depicted in panel (A).

